# The role of alternative splicing in marine-freshwater divergence in threespine stickleback

**DOI:** 10.1101/2024.06.14.598968

**Authors:** Carlos E. Rodríguez-Ramírez, Catherine L. Peichel

**Affiliations:** Division of Evolutionary Ecology, Institute of Ecology and Evolution, University of Bern, Bern 3012, Switzerland

**Keywords:** adaptation, alternative splicing, threespine stickleback, regulatory evolution, gene expression, gill

## Abstract

Alternative splicing (AS) regulates which parts of a gene are kept in the messenger RNA and has long been appreciated as a mechanism to increase the diversity of the proteome within eukaryotic species. There is a growing body of evidence that AS might also play an important role in adaptive evolution. However, the overall contribution of AS to phenotypic evolution and adaptation is still unknown. In this study we asked whether AS played a role in adaptation to divergent marine and freshwater habitats in threespine stickleback (*Gasterosteus aculeatus*). Using two published gill RNAseq datasets, we identified differentially expressed and differentially spliced genes (DEGs and DSGs) between population pairs of marine-freshwater stickleback in the Northeast Pacific and tested whether they are preferentially found in regions of the genome involved in freshwater-marine divergence. We found over one hundred DSGs, and they were found more often than expected by chance in peaks of genetic divergence and quantitative trait loci (QTL) that underlie phenotypic divergence between ecotypes. The enrichment of DSGs in these regions is similar to the enrichment of DEGs. Furthermore, we find that among the different types of AS, mutually exclusive exon splicing is the most strongly correlated with genetic divergence between ecotypes. Taken together, our results suggests that AS might have played a role in the adaptive divergence of marine and freshwater sticklebacks and that some types of AS might contribute more than others to adaptation.

## Introduction

Gene regulatory evolution has long been proposed to be an important driver of phenotypic evolution (King and Wilson 1975). In particular, cis-regulatory evolution has been argued to be an important driver of phenotypic evolution due to its potential to reduce antagonistic pleiotropy compared to protein evolution (Carroll 2005; Hoekstra and Coyne 2007; Stern and Orgogozo 2008; Wittkopp and Kalay 2012; Bomblies and Peichel 2022). Supporting these ideas, changes in gene expression mediated by cis-regulatory mutations have been shown to underlie phenotypic evolution and adaptation to different environments (Stern and Orgogozo 2008; Rebeiz et al. 2009; Chan et al. 2010; Wittkopp and Kalay 2012; Indjeian et al. 2016; Hill et al. 2021; Wooldridge et al. 2022). However, the literature on the role of regulatory evolution in phenotypic evolution and adaptation has for the most part focused solely on changes in gene expression and until recently has ignored other mechanisms of gene regulation, such as alternative splicing, which is common in many eukaryotic lineages (Chen et al. 2014). When alternative splicing occurs, different combinations of exons, and sometimes introns are included in the mature mRNA, leading to alternative mRNA isoforms that might result in the translation of different proteins from the same gene (Bush et al. 2017; Singh and Ahi 2022; Verta and Jacobs 2022; Wright et al. 2022). There are five types of alternative splicing (AS) events: 1) exon skipping (ES), when an alternative exon is not included in the mRNA; 2) mutually exclusive exons (MXE), when one exon out of a group is always included in the mRNA, but never more than one of the exons at the same time; 3) intron retention (IR), when an intron is kept in the mRNA instead of being spliced out as usual; 4) alternative 3’ splice sites (A3SS); and 5) alternative 5’ splice sites (A5SS), when part of the 3’ or 5’ end of an exon is spliced out of the mRNA (Figure 1). One gene can undergo a combination of these types of alternative splicing events, allowing for multiple different mRNA isoforms of a gene and leading to an overall increase in the diversity of the proteome. This might be the reason why AS has been found to be correlated with complexity (as measured by cell-type diversity) across eukaryotic lineages (Chen et al. 2014).

**Figure 1.**
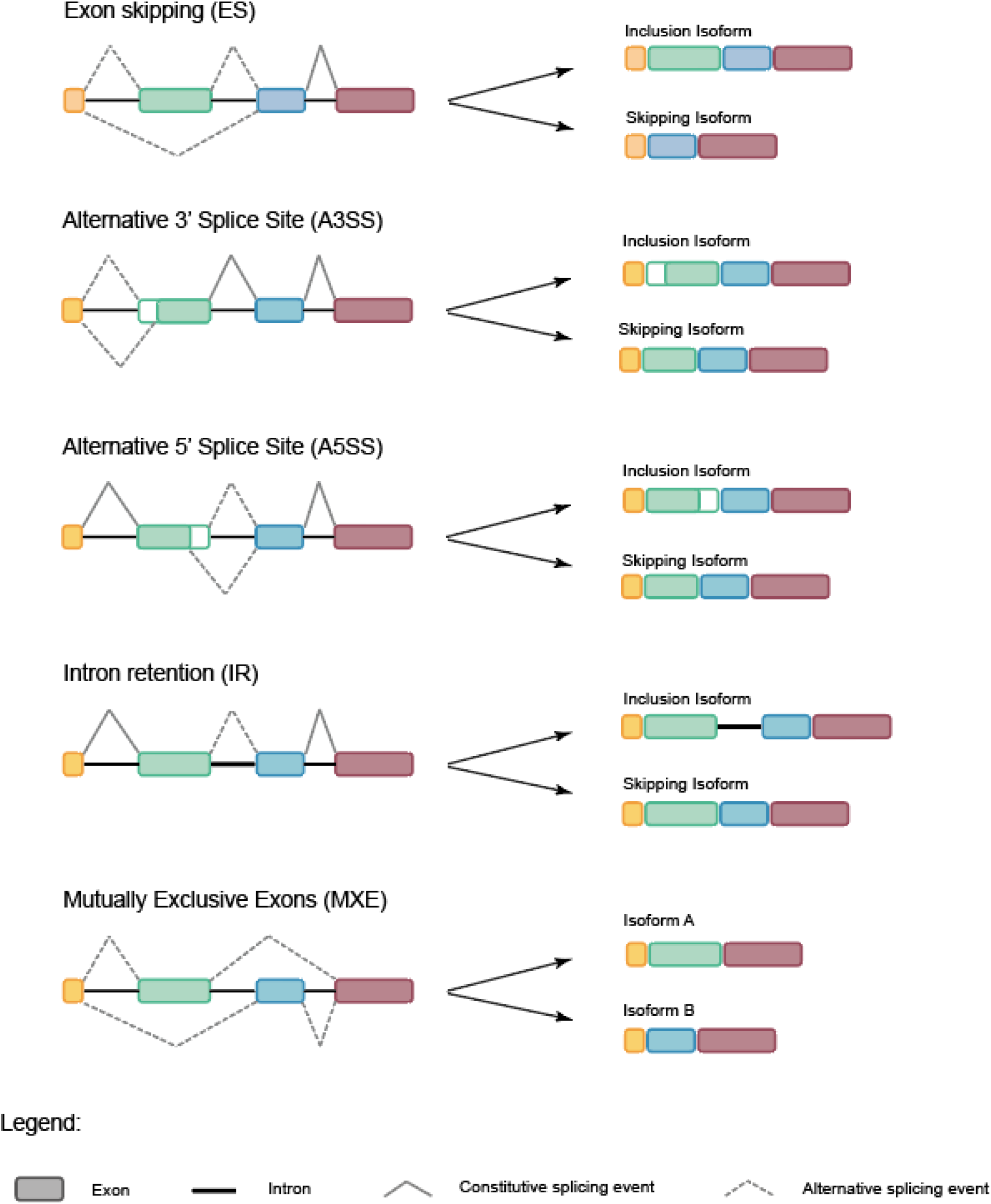
Different types of alternative splicing events. The DNA sequences are indicated to the left of the arrows, and the resulting mRNA sequences are indicated to the right. rMATs classifies one isoform as the “inclusion isoform” and another as the “skipping isoform”, then it counts all unambiguous reads mapping to each isoform to calculate the isoform inclusion difference (i.e change in the proportion of inclusion isoforms relative to skipping isoforms between ecotypes). For MXE splicing, the “inclusion” label is given to the isoform with the 5’-most exon.

The potential of AS to create different proteins has led to the question of whether it might play a role in phenotypic evolution and adaptation (Bush et al. 2017; Singh and Ahi 2022; Verta and Jacobs 2022; Wright et al. 2022). Indeed, recent studies have found that changes in AS can lead to phenotypic differences between ecotypes or species. For example, dorsal spine reduction in freshwater populations of threespine stickleback (*Gasterosteus aculeatus)* is mediated by the use of an A5SS in the first exon of *Msx2a,* a gene involved in osteoblast differentiation (Howes et al. 2017). Infrared sensation in vampire bats is linked to changes in the ratio of ES in the temperature receptor gene *Trpv1,* making it sensitive to temperatures around 30 ⁰C instead of 40 ⁰C (Gracheva et al. 2011). Increased lipid accumulation in two populations of the cavefish *Astyanax mexicanus* is caused by ES, resulting in a premature stop codon in the gene *per2,* a suppressor of lipid metabolism (Xiong et al. 2022). Eighteen other examples of intra or interspecific phenotypic variation mediated by alternative splicing are found at the genotype-phenotype database GePheBase (Martin and Orgogozo 2013; Courtier-Orgogozo et al. 2020), highlighting the potential of alternative splicing to cause phenotypic evolution. Supporting a role for AS in adaptation, many differentially spliced genes (DSGs) have been found between species or divergent ecotypes in human lice, cichlids, charr, sunflower and *Arabidopsis* (Tovar-Corona et al. 2015; Singh et al. 2017; Smith et al. 2018; Wang et al. 2019; Jacobs and Elmer 2021), and across environmental clines in wild house mice (Manahan and Nachman 2024). However, to our knowledge, to date only one study in benthic and pelagic ecotypes of Artic charr (Jacobs and Elmer 2021) has looked for evidence of divergent selection in DSGs between ecotypes of a species.

The threespine stickleback (*G. aculeatus*) is a good model to study questions related to the genetic basis of phenotypic evolution and adaptation. This small teleost fish is distributed across the Northern Hemisphere and the ancestral marine form independently colonised and adapted to many freshwater habitats approximately 12 000 years ago after the Last Glacial Maximum. Marine and freshwater stickleback diverge in ecology, physiology, and morphology, with the repeated evolution of many phenotypes in freshwater (Bell and Foster 1994). Genetic studies have found several of these parallel phenotypes are due to mutations in the same gene (Colosimo et al. 2005; Miller et al. 2007; Chan et al. 2010; Ishikawa et al. 2019). Many quantitative trait loci (QTL) mapping studies have identified regions of the genome strongly associated with phenotypic divergence (Peichel and Marques 2017). In addition, global genomic studies incorporating marine and freshwater population pairs from across the Northern Hemisphere have identified parallel peaks of genetic divergence across marine-freshwater population pairs that are putatively under divergent selection. Interestingly, most of these peaks of divergence are found in non-coding regions, hinting an important role of cis-regulatory evolution in the divergence between marine and freshwater sticklebacks. (Jones et al. 2012; Roberts Kingman et al. 2021).

Here, we use this system to ask whether alternative splicing could be a regulatory mechanism that contributes to adaptation to divergent environments. More precisely, we ask whether alternative splicing is important for marine-freshwater divergence in threespine stickleback. Using publicly-available RNA-seq data from marine and freshwater populations from the Northeast Pacific, we ask the following questions: 1) are there differentially spliced genes (DSGs) between the two ecotypes?; 2) is there evidence that these DSGs might mediate phenotypic divergence between the ecotypes?; and 3) is there any evidence that natural selection acted on these DSGs?

## Results

### Over one hundred DSGs between marine and freshwater stickleback

Using an RNA-seq dataset from gill tissue of marine and freshwater sticklebacks from Canada (Supplementary Table S1), we found 16 999 expressed genes, of which 1882 are differentially expressed genes (DEGs) between marine and freshwater samples. We detected alternative splicing events in 1345 genes, of which 139 are differentially spliced genes (DSGs) between ecotypes (Figure 2a, Supplementary Table S2). Thirty genes are simultaneously DEGs and DSGs (differentially expressed and spliced genes, or DESGs). We found all five types of differential splicing events amongst the DSGs (Figure 2b). Differential MXE is the most common and is present in 71 DSGs; this is followed by 45 DSGs with differential ES, 22 DSGs with IR, 18 DSGs with A3SS, and 8 DSGs with differential A5SS (Figure 2b, Supplementary Table S2). There is almost no enrichment of GO Terms in DEGs or DSGs. The only exception is an enrichment for the terms “phosphatidylinositol monophosphate phosphatase activity” and “phosphatidylinositol-3-phosphate phosphatase activity” in the DEGs (Supplementary Table S3).

**Figure 2.**
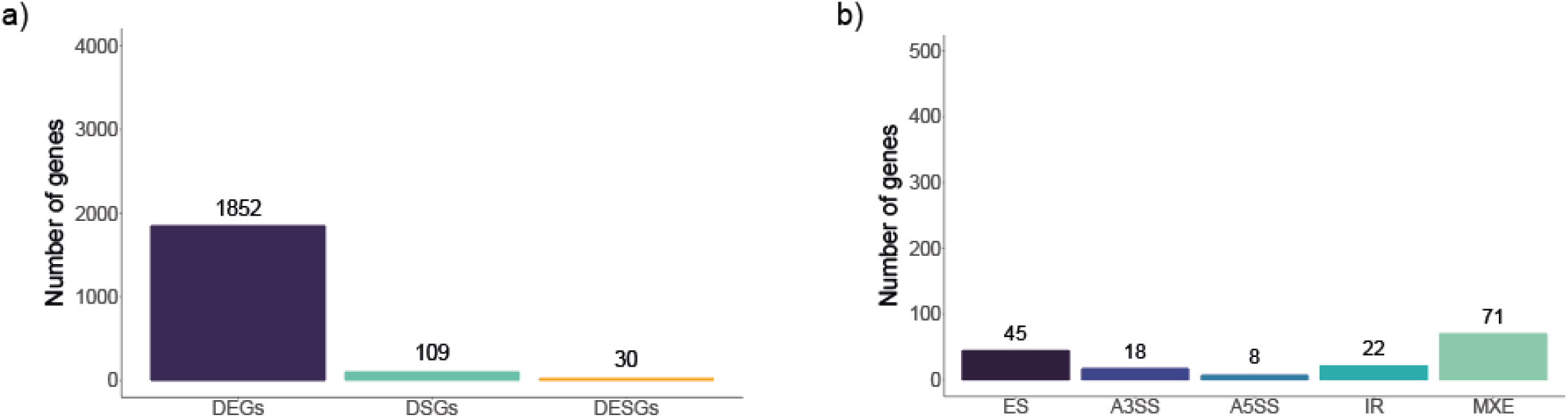
a) Number of differentially expressed genes (DEGs), differentially spliced genes (DSGs), and differentially expressed and spliced genes (DESGs) in the gill transcriptome. DEGs and DSGs counts in the figure do not include DESGs. b) Number of differentially spliced events of each AS type found within the 139 DSGs (including DESGs) in the gill transcriptome.

### Pacific DSGs are significantly enriched in some categories of QTL

To test whether DEGs and DSGs might be mediating phenotypic divergence between marine and freshwater sticklebacks, we tested their enrichment in 316 QTL that underlie traits that diverge between Pacific marine and freshwater populations (Supplementary Table S4) (Peichel and Marques 2017; Liu et al. 2022; Rennison and Peichel 2022). The QTL span most of the gill transcriptome: out of the 16 999 genes in this dataset, 12 129 genes (71.4%) are inside at least one QTL. We found that DEGs are enriched in the overall set of QTL, as well as in most phenotypic sub-categories (Figure 3 and Supplementary Table S5). Meanwhile, the DSGs are not enriched in the overall set of QTL, but they are enriched in QTL sub-categories associated with body shape, defence and feeding (p-value = 0.001, 0.013 and 0.033, respectively; permutation tests, 1000 permutations). DESGs are also enriched in QTL associated with body shape and swimming (p-value = 0.01 and 0.009, respectively; permutation tests, 1000 permutations) (Figure 3 and Supplementary Table S5).

**Figure 3.**
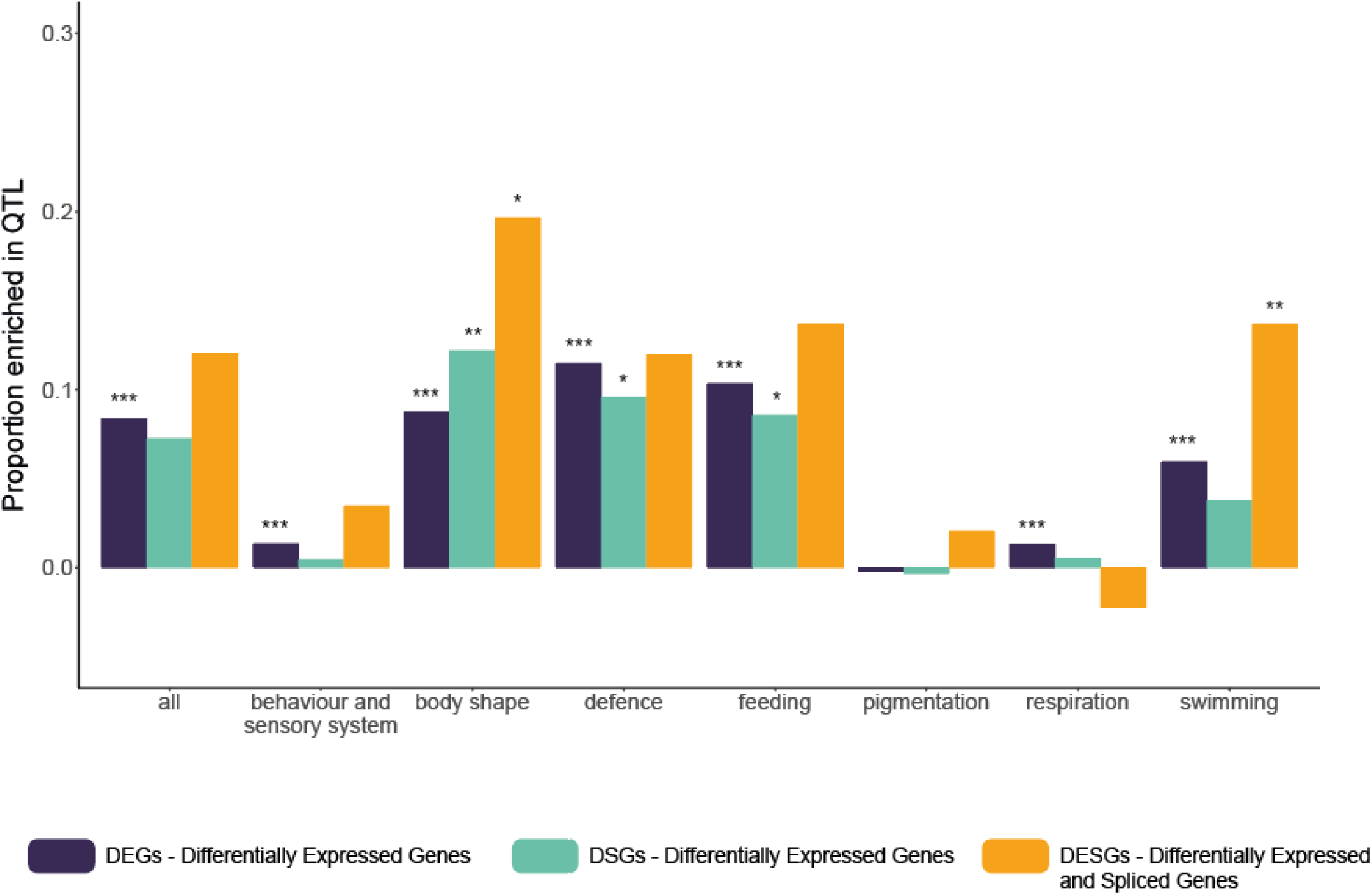
QTL enrichment analysis results for DEGs (dark blue), DSGs (aqua), and DESGs (orange). Bars represent the proportion of DEGs, DSGs, or DESGs in a QTL for a given category minus the proportion of background genes in a QTL for that category. Asterisks represent significance levels for the QTL enrichment test (permutation test, 1000 permutations): * p-value < 0.05; ** p-value < 0.01; *** p-value < 0.001.

### DEGs and DSGs are significantly enriched in EcoPeaks

To determine whether DEGs and DSGs might be under divergent selection, we tested whether they are significantly enriched in EcoPeaks, which are regions of the genome with peaks of genetic divergence between multiple marine and freshwater populations from either the Northeast Pacific (“Pacific EcoPeaks”) or from the Northeast Pacific, California, and Europe (“Global EcoPeaks”) (Roberts Kingman et al. 2021). The Pacific EcoPeaks include 22.7% of the genes in our transcriptome (3854 out of 16 999) (Supplementary Table S2). We found 43.4% of DEGs (803 out of 1852), 40.4% of the DSGs (44 out of 109) and 56.7% of the DESGs (17 out of 30) are inside the Pacific EcoPeaks. This is significantly more than the 22.7% to 23.7% of background genes that are in these regions of the genome (Figure 4a, Supplementary Table S6). DEGs and DSGs are similarly enriched in Global EcoPeaks (Figure 4b, Supplementary Table S6).

**Figure 4.**
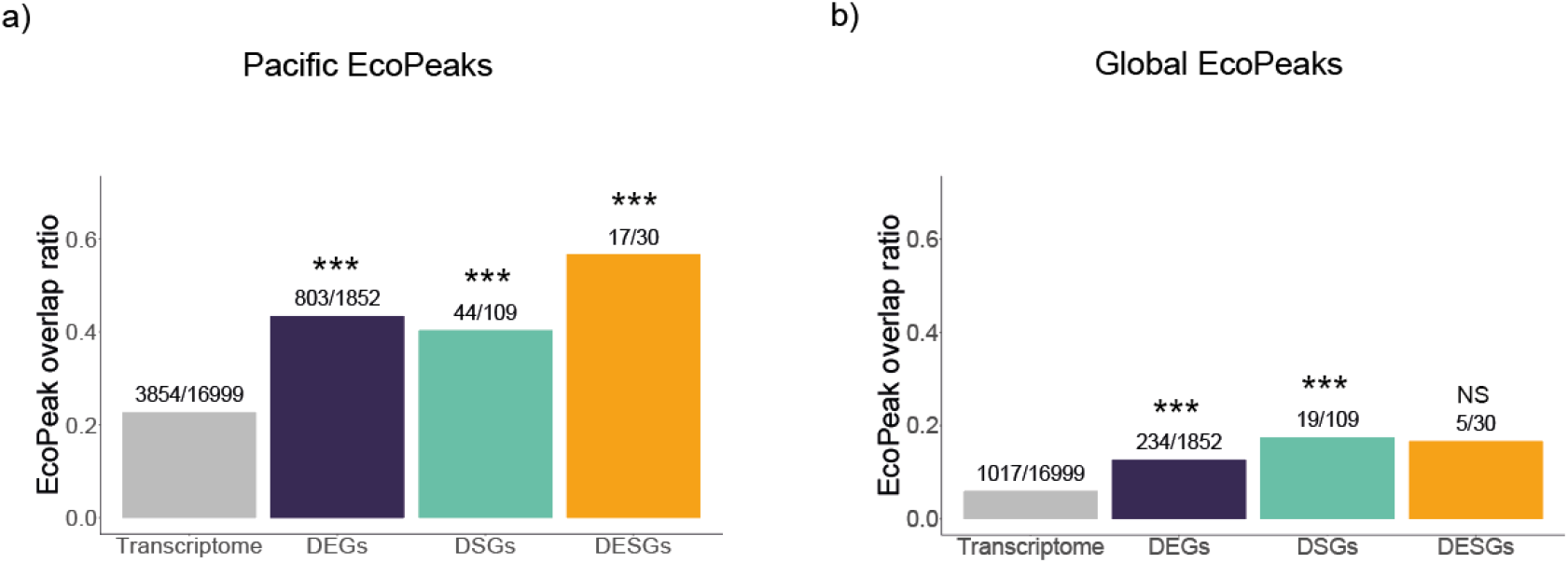
Proportion of DEGs, DSGs, DESGs and transcriptome genes overlapping Pacific (a) and Global (b) EcoPeaks. Transcriptome genes are all genes that passed the minimum expression filter (i.e the background of the DEGs). DSGs and DESGs were compared to their respective backgrounds, but for simplicity they are not represented in the figure (Supplementary Table S6). The number of genes of each category inside the EcoPeaks relative to the total number of genes in that category are shown. Asterisks represent significance levels for the EcoPeak enrichment test (permutation test, 1000 permutations): * p-value < 0.05; ** p-value < 0.01; *** p-value < 0.001; NS – not significant, p-value > 0.05.

DEGs and DSGs overlapping both EcoPeaks and QTL are the strongest candidates for genes with an adaptive role in the marine-freshwater divergence in threespine stickleback. Thus, we asked whether DEGs and DSGs overlapping Pacific EcoPeaks and QTL are enriched in any particular biological functions. However, we found that these genes are not significantly enriched in any GO Term category (Supplementary Table S3). Thus, we used the GeneCards database to investigate the function of the 15 DSGs found within both Global EcoPeaks and QTL. We found multiple genes involved in essential amino acid metabolism, chromatin remodelling, immunity, vesicle transport and muscle function (Supplementary Table S7).

Examining the types of differential splicing events underlying the DSGs, we found that DSGs with significant MXE, ES and IR events are enriched in Pacific EcoPeaks (54.9%, 37.8% and 31.8% respectively; p-values < 0.001, 0.021 and 0.034 respectively; permutation test, 1000 permutations) (Supplementary Figure S1a, Supplementary Table S6). Though not significant, DSGs with A3SS show a similar trend, with 7 out of 18 (38%) found in EcoPeaks (p-value = 0.172; permutation test, 1000 permutations). DSGs with MXE and IR are also significantly enriched in the Global EcoPeaks (Supplementary Figure S1b, Supplementary Table S6).

Finally, we also examined whether there is a difference in the overall fold-change in expression (DEGs) or the inclusion isoform change (“isoform difference” in DSGs) between genes inside versus outside of EcoPeaks. We found that DSGs in both Pacific and Global EcoPeaks have a significantly higher isoform difference than those outside, particularly DSGs within Global EcoPeaks (Figure 5, Table S8). Interestingly, DEGs do not differ in their fold-change inside and outside of EcoPeaks (Figure 5, Table S8).

**Figure 5.**
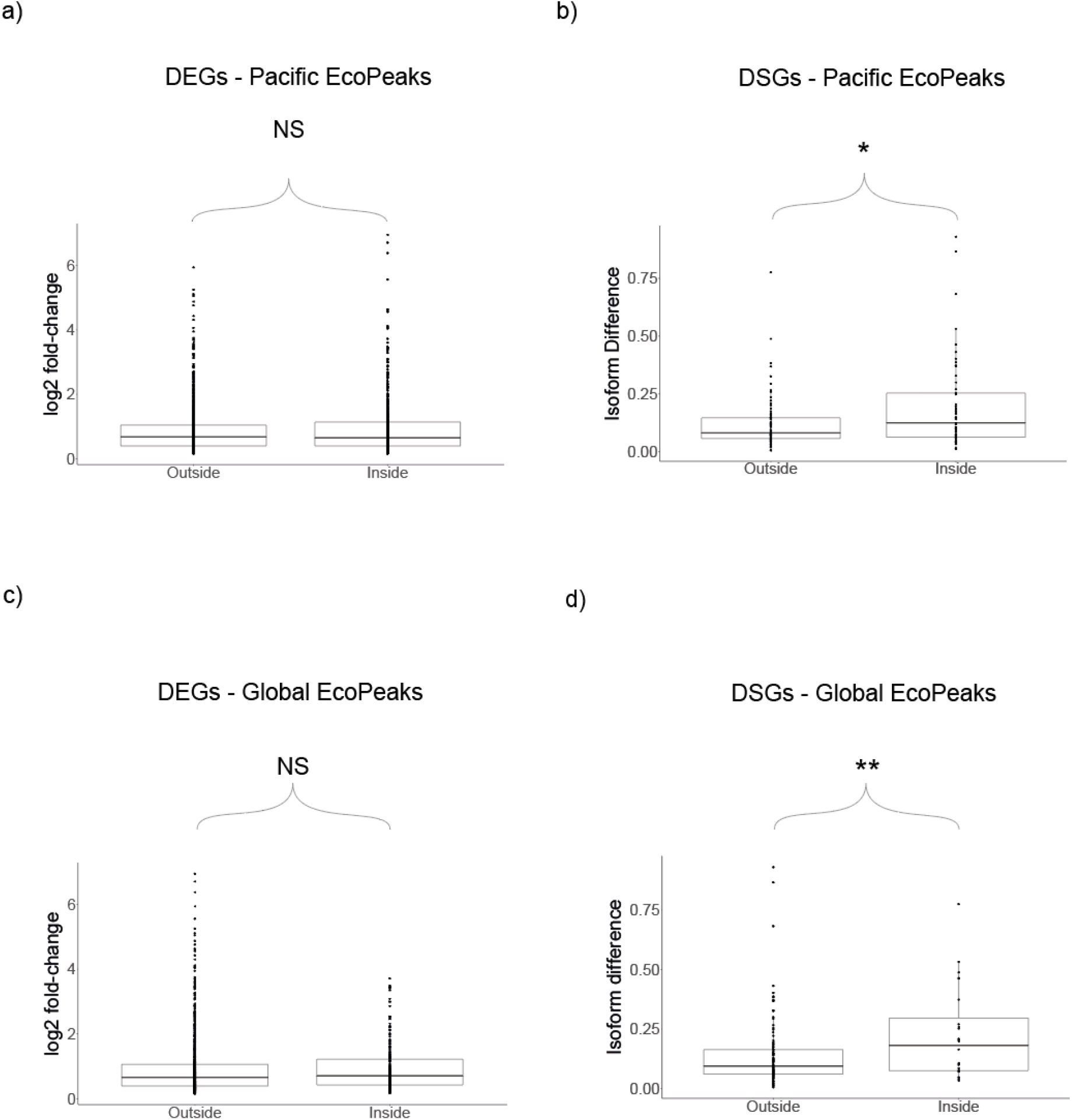
Difference in fold-change (a and c) and isoform difference (b and d) of DEGs and DSGs inside and outside of Pacific (a and b) or Global (c and d) EcoPeaks. Asterisks represent significance levels for the difference of the medians (permutation test, 1000 permutations): * p-value < 0.05; ** p-value < 0.01; NS – not significant, p-value > 0.05.

### No strong correlation between transcriptome-wide genetic divergence and differential expression and splicing

To determine to what extent marine-freshwater splicing divergence is correlated with marine-freshwater genetic divergence, we compared the isoform difference of all 1345 genes used in the differential splicing analysis with their average SNP p-value from the Pacific EcoPeak data (Roberts Kingman et al. 2021). We found a very weak positive correlation between isoform difference and genetic distance (R2 = 0.015, p-value = 1.3e-06) (Supplementary Figure S2). We found a similarly weak positive correlation between genetic distance and the log of expression fold-change between ecotypes (R2 = 0.010, p-value = 2.2e-16) (Supplementary Figure S2). These weak correlations are likely driven by the EcoPeaks. When we separated the data inside and outside of the EcoPeaks, the correlations between genetic distance and isoform difference/fold-change are weaker, and in most cases no longer significant (Supplementary Figure S2).

### Differences in effect size and genetic divergence between types of AS

To gain insights into whether certain types of splicing could be more important to adaptation than others, we tested whether genetic divergence and strength of DS changes between the five types of AS events. For this analysis, we classified DSGs by the type of AS event that was the strongest DS event in that gene. When comparing the different types of AS with each other, there is a tendency for genes with MXE and IR to have a higher isoform difference between ecotypes, though this is only significant in the comparison between MXE and ES (Supplementary Figure S3a and Table S9a). Regarding genetic divergence, there is a tendency for genes with MXE to be more divergent than other types of DSGs, but it is only significant when compared to IR (Supplementary Figure S3b and Table S9b). However, when we compare the DSGs to their non-DSG counterparts for each type of AS, we find that DSGs with differential MXE, ES and IR have a significantly higher genetic divergence than genes that have MXE, ES and IR, respectively, but that are not differentially spliced between ecotypes (Figure 6 and Table S10). This suggests that DSGs with these types of splicing are more likely to have been targeted by selection between marine and freshwater sticklebacks.

**Figure 6.**
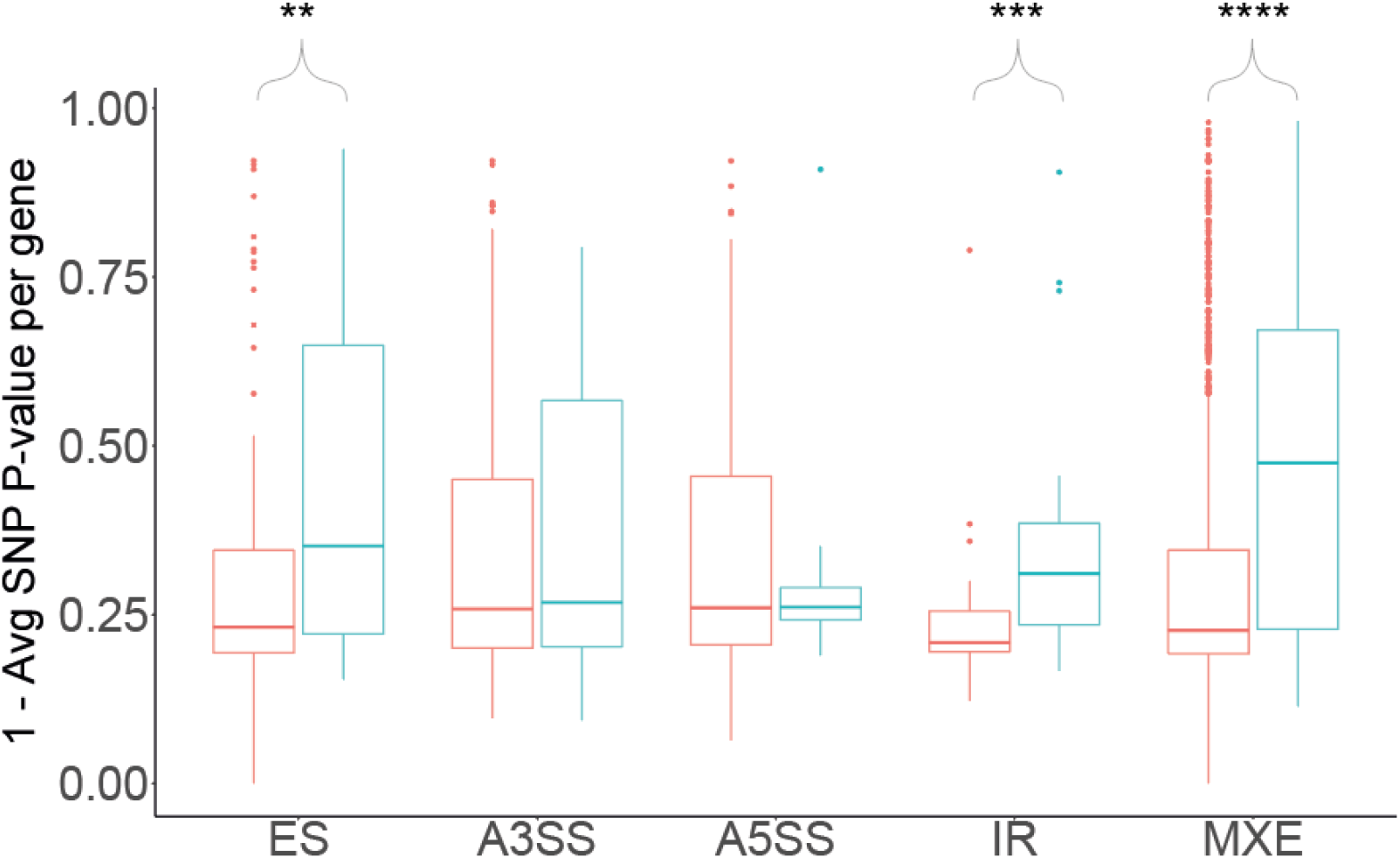
Distributions of average SNP p-value for significant DSGs (blue) and non-DSGs (red) per splicing type. Non-DSGs are genes that are alternatively spliced but not significantly differentially spliced between marine and freshwater samples. To make the Y axis more intuitive, the average SNP p-value is subtracted from 1 so that larger values represent greater genetic divergence. Asterisks indicate whether DSGs have a significantly higher genetic divergence median than non-DSGs (permutation test, 10,000 permutations): * p-value < 0.05; ** p-value < 0.01; *** p-value < 0.001; **** p-value < 0.0001.

## Discussion

The magnitude of the role of alternative splicing in adaptation is still unknown. We sought to tackle this question by assessing the role of alternative splicing in the marine-freshwater divergence of threespine stickleback (*G. aculeatus*). We show that there are more than one hundred DSGs in the gill transcriptome between marine and freshwater stickleback populations from the Northeast Pacific. DSGs are enriched not only in regions under putative divergent selection in the genome, but also within QTL underlying phenotypic divergence between ecotypes in the Pacific. Furthermore, the enrichment of DSGs in these regions of the genome was similar to the enrichment found for DEGs, suggesting that DSGs might be as important for adaptation as DEGs. Among the five types of AS, MXE splicing events are the most commonly divergent between ecotypes and are in the most genetically divergent genes. Thus, mutually exclusive exon use could be particularly important for adaptation to different environments. Taken together our results suggest that alternative splicing might play an important role in marine-freshwater divergence in threespine stickleback.

### Over one hundred genes have changes in alternative splicing between marine and freshwater sticklebacks in gill

To determine the role of alternative splicing in the freshwater-marine divergence in threespine stickleback, we first assessed the extent of differential expression and differential splicing between ecotypes. We found 1852 DEGs, 109 DSGs and 30 DESGs (Figure 2a). This disparity in abundance of DEGs and DSGs is a pattern commonly found in other studies (Grantham and Brisson 2018; Jacobs and Elmer 2021; Steward et al. 2022; Rodríguez-Ramírez et al. 2023), though not always (Singh et al. 2017). This might suggest that differential splicing is less commonly used than differential expression for adaptation. Since AS directly affects the protein sequence it is possibly under stronger purifying selection than differential expression. However, it is important to note that methods to detect AS in short-read data have greater limitations in their ability to identify and quantify changes in splicing than methods for identification of DEGs in short-read data. For example, methods based on splice junctions like the one used in this study and others (e.g. Steward et al., 2022) work mainly with reads overlapping these junctions. Because most reads cannot be used for analyses, the power of the DS analyses is greatly reduced compared to the DE analyses, which can use most reads in the transcriptome. Methods based on differential exon usage have been used in other studies (e.g. Singh et al., 2017; Jacobs & Elmer 2021; Rodríguez-Ramírez et al., 2023) and can use most of the same reads as DE analyses but cannot detect certain types of splicing (i.e. A3SS and A5SS). Thus, all AS studies based on short-read RNA-seq data are likely to underestimate the extent of DS. In contrast, long-read RNA sequencing methods like Iso-seq, give us information on the full mRNA sequence and have revealed many previously unknown isoforms both in animals and plants (Singh and Ahi 2022). Future studies using long-read RNA sequencing technologies are essential to properly assess the relative role of DEGs and DSGs in adaptation.

### DSGs are associated with regions underlying phenotypic divergence in body shape, defence and feeding between marine and freshwater stickleback

Consistent with a role for DEGs and DSGs in mediating phenotypic divergence between marine and freshwater stickleback, we found that DEGs and DSGs were enriched in QTL associated with a variety of phenotypes. While DEGs were enriched in more QTL, DSGs were still associated with QTL underlying phenotypes like body shape, defence traits and feeding traits. However, an important caveat is that these QTL are mostly associated with phenotypes completely unrelated to gill tissue, except for some of the feeding trait QTL (Supplementary Table S4). The expression of genes associated with these QTL in gill tissue could be due to several non-mutually exclusive reasons. One possibility is that some of these genes have pleiotropic effects in both gill and the tissues associated with the QTL. For example, the gene *stat3* is amongst the most divergent DSGs in the dataset (Supplementary Table S7) and is a pleiotropic transcription factor with described roles in multiple biological processes including skeletal development, hair maintenance, immunity and cellular respiration (Levy and Lee 2002; O’Shea et al. 2002; Wegrzyn et al. 2009; Hillmer et al. 2016; Zhou et al. 2021). Furthermore, *stat3* in mice is known to produce two alternative splicing isoforms: a full-length isoform, *stat3α;* and a truncated isoform, *stat3β.* The isoforms have different phenotypic effects and are thought to partially explain the high pleiotropy of this gene (Maritano et al. 2004). A second possibility is that most of the QTL have very low resolution and span large regions of the genome, so most of the genes found within them are unlikely to be related to the focal QTL phenotype. An enrichment of DEGs and DSGs in these QTL could occur if these genes affect other unmapped phenotypes that are linked to these QTL, consistent with the QTL clustering observed in stickleback (Peichel and Marques 2017). A final possibility is that strong selection for DE and DS in a gene within a specific tissue could allow the fixation of regulatory variants that cause leaky expression and splicing in unrelated tissues like gill. Mutations in cis-regulatory elements are known to affect gene expression noise (Richard and Yvert 2014) and the strength of selection against noise depends on the function of the gene and its position in the gene pathway (Barroso et al. 2018). Thus, it is possible that when rapid adaptation to different environments occurs, regulatory variants that are favourable in one tissue might get fixed despite increasing the transcriptional noise of that gene in other tissues. While the DEGs and DSGs that we found are enriched in QTL and EcoPeaks between marine and freshwater sticklebacks, many of the DEGs and DSGs have relatively small effect sizes (Supplementary Table S2), which is what we would expect if some of these genes represented some sort of transcriptional leakage resulting from divergent selection between marine and freshwater stickleback in other tissues.

### DSGs are associated with regions under divergent selection between ectoypes

Having determined that DSGs are enriched in regions of the genome associated to phenotypic divergence between marine and freshwater sticklebacks, next we asked whether the same was true for the smaller subset of the genome with putative signatures of divergent selection between ecotypes. We used the EcoPeaks database (Roberts Kingman et al. 2021) and found that DEGs and DSGs were enriched in both Pacific and Global EcoPeaks (Figure 4). This suggests that DSGs are as likely as DEGs to be under divergent selection between threespine stickleback ecotypes in Northeast Pacific populations. It is possible that some of these genes are DE and DS simply because their causative cis-regulatory variants merely hitchhiked with the actual targets of selection in the EcoPeaks. However, we find only a very weak linear correlation between genetic divergence in the EcoPeaks data and the strength of DS and DE across the gill transcriptome (Supplementary Figure S2), suggesting that the DEGs and DSGs inside of the Pacific EcoPeaks are not just a side-effect of local genetic divergence.

There is not a strong linear correlation between genetic distance in the Pacific EcoPeak data and the strength of DE or DS in the gill transcriptomes. However, it is important to note that we do not have genomic sequencing data for the individuals used to identify DEGs and DSGs, so we cannot directly look at the correlation between genetic distance and DE and DS. Nonetheless, we do find that DSGs in Global EcoPeaks have stronger DS than those outside of Global EcoPeaks (Figure 5). We find a similar trend for DSGs in the Pacific EcoPeaks (p-value = 0.059, Figure 5, Supplementary Table S8). Interestingly, we do not find an increase in the fold-change of DEGs inside and outside of EcoPeaks (Figure 5, Supplementary Table S8). If the strength of DE and DS is correlated with their adaptive importance, this result suggests selection for stronger splicing divergence in the EcoPeaks but not for stronger expression divergence. It is also possible that divergent selection is equally strong in DE and DS, but splicing-mediated phenotypic effects require greater splicing divergence than expression-mediated phenotypic effects. Finally, it is possible that DS is less constrained by purifying selection and thus diverges more quickly inside the EcoPeaks, potentially through hitchhiking of splicing regulatory variants with other targets of selection. However, if this hypothesis were true, we would expect to find a correlation between genetic divergence and DS inside and outside of the Pacific EcoPeaks, which we do not (Supplementary Figure S2). Taken together, our results suggest that there is selection for greater splicing divergence than expression divergence between marine and freshwater sticklebacks inside the EcoPeaks.

### MXE, IR and ES may play an important role in marine-freshwater divergence

Alternative splicing can generate different types of splicing events (Figure 1). To test whether they might play different roles in adaptation, we assessed the presence of five types of AS in our data. We found all five types of AS in the DSGs of our dataset. MXE, ES and IR events are not only more common, but they are also the only types of AS enriched in EcoPeaks (Supplementary Figure S1). Furthermore, we also find that only DSGs with either of these three types of differential AS events have a higher genetic divergence their non-DSGs counterparts. Taken together these results suggest that MXE, ES and IR are more likely than A3SS and A5SS to be under selection between the marine-freshwater sticklebacks and play a role in their adaptive divergence.

It is not clear why A3SS and A5SS would be less used in adaptation. Just like ES and IR, A3SS and A5SS remove and add mRNA sequence and thus have the potential to increase proteomic diversity. However, most of the time this likely leads to aberrant proteins, truncated proteins or degradation of the mRNA by nonsense-mediated mRNA decay (NMD). Nonetheless, all of these can still be a mechanism for indirect down-regulation of genes, which could be adaptive. This is the case for gene *Msx2a*, which underlies dorsal spine differences between marine and freshwater threespine sticklebacks. The freshwater allele of *Msx2a* leads to shorter dorsal spines and has increased A5SS splicing of the first exon of this gene, which introduces an early stop codon and leads to a truncated non-functional version of the protein (Howes et al. 2017). Similarly, an ES event in the gene *TBXT* removes part of the transcriptional regulation domain of the protein and leads to the loss of tail in apes (Xia et al. 2024). Overall, ES and IR probably tend to add or remove larger sequences to the mRNA than A3SS and A5SS, which could increase the probability of new protein domains to emerge. This might also make down-regulation of genes by NMD more reliable through ES and IR than A3SS and A5SS. Indeed, IR has been found to both regulate gene expression by NMD and to contribute to functional isoforms (Wong et al. 2016). ES has similarly been found to regulate the incorporation of “poison” exons in mammalian splicing regulator (SR family) genes that lead to NDM but are important for cell fitness (Wright et al. 2022).

Furthermore, ES can also regulate new exons that emerge in intronic sequences. This can occur for example, through exonization of intronic transposable elements (TEs) (Lev-Maor et al. 2007; Sorek 2007; Wright et al. 2022). Studies in human and mice suggest that most new exons are alternatively spliced and expressed at low frequencies, making them nearly neutral (Xing and Lee 2006; Sorek 2007). This can allow new exons to evolve and potentially acquire a new function over time. For example, primates possess many lineage-specific exons that have originated from a class of TEs known as *Alu* elements. While their function is still not clear, *Alu*-containing exons can be expressed. Although they tend to be alternatively skipped (Sorek et al. 2002), some have been found to contribute to the sequence of proteins (Lin et al. 2016; Martinez-Gomez et al. 2020; Wright et al. 2022). While over 79% of new exons in humans are predicted to be deleterious (Sorek 2007), some are known to have acquired new functions. For example, the human ADAR2 gene is an RNA editing enzyme with a primate-specific exon 8 derived from an *Alu* element. This exon can be included in the catalytic region of the protein, which still works with the same substrate but with an altered catalytic activity (Sorek 2007).

Despite the potential of ES and IR to contribute to adaptation, MXE is the type of splicing that stood out the most in our data. This is the most common differential AS event (Figure 2), the most enriched in EcoPeaks (Supplementary Figure S1), and the AS event found in the DSGs with the greatest isoform differences and genetic divergence (Figure 6; Supplementary Figure S3). MXE is a type of splicing that switches one alternative exon for another in the mRNA (Figure 1). This allows for modular changes in the protein structure where one protein domain can be switched for a different one without affecting the integrity of the protein. One extreme example of this is the *Dscam* gene in *Drosophila melanogaster,* which encodes for a transmembrane protein that is important for neuronal connection and self-avoidance. The gene has 115 exons, of which 95 belong to one of four clusters of mutually exclusive exons and has been found to produce up to 18 496 isoforms (Sun et al. 2013), more than the number of genes in the genome of *D. melanogaster.* Furthermore, MXE is thought to be associated with exon duplications (Kondrashov 2001; Letunic 2002; Wright et al. 2022). Similar to gene duplications and exonization of intronic sequences, MXE can allow for the evolution of new functions in one of the paralogous exons while maintaining its ancestral function in the other paralog. Evolving new exons through duplications rather than exonization of TEs could be advantageous in that duplicated exons are immediately able to code for a functioning part of the existing protein. Thus, exon duplication and combined with MXE could promote the evolution of alternative protein domains that might become advantageous when the species adapts to a different environment. Taken together, our results suggest that MXE could be a powerful mechanism to maintain standing genetic variation in protein isoforms and could play an important role in the divergence of marine and freshwater stickleback in the Pacific.

## Conclusion

The role of alternative splicing in adaptation is still poorly understood. However, alternative splicing can be quite a versatile regulatory mechanism that can both indirectly down-regulate the expression of a gene by creating non-functional isoforms and mediate modular changes in the protein by permuting different exons from the gene into the mRNA. Consistent with a growing body of evidence in other systems, we find evidence for a role of alternative splicing in the marine-freshwater adaptive divergence of threespine stickleback. DSGs between marine and freshwater populations are enriched in QTL underlying phenotypic divergence between ecotypes, and DSGs are as enriched as DEGs in regions of the genome putatively under divergent selection between marine and freshwater populations. We also find evidence that different types of alternative splicing might contribute differently to adaptation, with MXE standing out in our data and suggesting that the modular change of exons could be particularly important for adaptation. Finally, our results are quite likely an underestimate of the true extent of differential splicing between marine and freshwater sticklebacks. Limitations of geographical representation, number of tissues, developmental time points and short-read data mean that we are probably detecting only a small part of the alternative splicing events in the threespine stickleback transcriptome. Future studies with more populations and tissues, long-read sequencing, and functional analyses will be essential to have a more precise picture of the role of alternative splicing in marine-freshwater divergence in threespine stickleback.

## Materials and Methods

### RNA-seq data

We searched the NCBI Sequence Read Archive (SRA) for publicly available RNA-seq data that met three criteria: 1) the data included samples from marine and freshwater population pairs of threespine stickleback (*G. aculeatus*); 2) the data came from the same tissue, so it could be merged and compared; 3) the data came from Pacific populations, since these are the ones with the most complete QTL and genetic divergence data (Peichel and Marques 2017; Roberts Kingman et al. 2021). Following these criteria, we found data from two RNA-seq studies in stickleback gill tissue (Gibbons et al. 2017; Verta and Jones 2019) (Supplementary Table S1). The Verta and Jones (2019) data is from first-generation descendants of wild-caught individuals grown in a common garden in the lab at 3.5 parts per thousand (ppt) salinity. From this dataset we used four marine and four freshwater individuals from the Little Campbell River (British Columbia, Canada). The Gibbons et al. (2017) data included ten marine wild-caught individuals from Oyster Lagoon (British Columbia, Canada) that spent four weeks at 20 ppt in the laboratory before being gradually moved to either 0.0 ppt (five individuals) or 30 ppt (five individuals) for three months, and ten freshwater wild-caught individuals from Trout Lake (British Columbia, Canada) that spent four weeks at 2.0 ppt in the laboratory before gradually being exposed to either 0.0 ppt (five individuals) or 30 ppt (five individuals) for three months.

### RNA-seq data pre-processing

Quality control of the RNAseq read libraries was done with FastQC v0.11.7 (https://www.bioinformatics.babraham.ac.uk/projects/fastqc/). Reads were quality trimmed using Trimmomatic v0.36 (Bolger et al. 2014). Reads where both paired ends passed quality filtering were then mapped against version 5 of the *G. aculeatus* genome (Nath et al. 2021) using STAR v2.7.10b (settings: --twopassMode --chimSegmentMin [1/3 of read length] --alignSJDBoverhangMin 3 -- alignIntronMin 70 --alignIntronMax 562000 --alignMatesGapMax 562000 --limitSjdbInsertNsj 2000000). Library quality metrics from FastQC and alignment quality metrics from STAR and featureCounts were summarized and visualized with MultiQC v1.14 (Ewels et al. 2016) to assess sample quality (Supplementary Table S1).

### Identification of differentially expressed genes (DEGs)

We obtained count tables of the RNA-seq reads mapped to each gene in the genome using the gene annotations from NCBI (build 100) for version 5 of the *G. aculeatus* genome (https://www.ncbi.nlm.nih.gov/datasets/taxonomy/481459/) and *featureCounts* from the *Subread* v2.0.3 (Liao et al. 2014) package. We excluded reads where one of the pairs was unaligned, or when the two pairs mapped to different chromosomes or different strands. Next, we used the *edgeR* v3.28.1 package (Chen et al. 2008) in *R* v3.6.1 (R Core Team 2019) for the differential expression analysis. First, we filtered lowly-expressed genes using the *filterByExpr()* function, which removes genes with less than 10 counts in a minimum number of samples based on sample size (for further details consult the *filterByExpr()* documentation). For our data, this meant that genes with less than ten counts in nine or more samples were removed. Second, we fit the count data in *edgeR* to a negative binomial general linear model to control for batch effects present in the datasets. We used a ∼(study+ecotype) model to control for the effect of having data from two different studies. Third, we ran a quasi-likelihood F test on the fitted data to test for differential expression between marine and freshwater samples.

### Identification of differentially spliced genes (DSGs)

To identify DSGs we used a method based on the identification of splicing events in RNA-seq data implemented in the program *rMATs* v4.1.2 (Shen et al. 2014). First, rMATs uses reads that STAR maps to exon boundaries in the mRNA or read pairs that map to different exons in a gene, to identify five types of alternative splicing events: Exon skipping (ES), alternative 5’ and 3’ start sites (A5SS and A3SS), intron retention (IR), and mutually exclusive exons (MXE). For each splicing event, rMATs defines an inclusion isoform and a skipping isoform (Figure 1), counts how many reads map to each isoform, and calculates an Isoform Inclusion Difference metric (Shen et al. 2014), also known as Percent Spliced-In, or PSI (Grantham and Brisson 2018; Rogers et al. 2021). This metric, which we will refer to as isoform difference, measures how much the ratio of the inclusion isoform changes between treatments (Shen et al. 2014). Since rMATs does not include batch-correction or minimum expression filters, we took the raw counts of the splicing events identified by *rMATs* and used the *edgeR* function *filterByExp(),* to filter out lowly expressed splicing events. Then, we corrected for the study effect in the data just as in the differential expression analysis using the function *ComBat-seq(),* from the package *sva* v3.35.2, which outputs batch-corrected counts (Zhang et al. 2020). Finally, using the “*—task stat*” mode in *rMATs,* we re-calculated the isoform differences using the batch-corrected counts from the splicing events and did the statistical test for differential splicing. We used the default rMATs settings for the analysis.

### EcoPeak and QTL enrichment analyses

To test whether DEGs and DSGs might be under divergent selection, we asked whether they were enriched in “EcoPeaks”, regions of the genome with peaks of genetic divergence between multiple marine and freshwater populations from across the Northern hemisphere (Roberts Kingman et al. 2021). The dataset is divided into Northeast Pacific (which we will refer to as Pacific for simplicity) and Global EcoPeaks depending on the samples used. The Pacific EcoPeaks were identified by comparing 12 marine and 57 freshwater populations from Alaska (US), Haida Gwaii (Canada), British Columbia (Canada), and Washington State (US). The Global EcoPeaks were identified by comparing 28 marine and 56 freshwater populations from the Northeast Pacific, California, and Europe. EcoPeaks were identified using two approaches: 1) a window-based genetic distance approach; and 2) a SNP-level statistical test for imbalance of genetic variants between marine and freshwater populations. Sensitive EcoPeaks were defined when an FDR of 5% was obtained in either of the two analyses, while Specific EcoPeaks were defined when an FDR of 1% was obtained in both analyses (for more details consult Roberts Kingman et al., 2021). Since the number of DSGs in our dataset was not very high, we used the Sensitive EcoPeaks to increase the power of our analyses. We tested the enrichment of our DEGs and DSGS in both Pacific and Global EcoPeaks. Since the EcoPeaks were originally identified in the v4.1 (*gasAcu1-4*) genome assembly (Roberts Kingman et al. 2021), we lifted over the coordinates of the EcoPeaks to the coordinates of the version 5 genome (Supplementary Table S11) using the *liftOver()* function of the *R* package *rtracklayer* v1.46.0 (Lawrence et al. 2009) and the chain file “v4.1_to_v5.chain” available at the Stickleback Genome Browser (https://stickleback.genetics.uga.edu/).

Briefly, the enrichment analysis involved: 1) identifying the proportion of DEGs and DSGs that overlapped with EcoPeaks; 2) identifying the proportion of background genes that overlapped with EcoPeaks; and 3) testing if the proportion of DEGs and DSGs in EcoPeaks is significantly higher than the proportion of background genes using a permutation test (1000 permutations) in R. The background genes included in this analysis corresponded to the genes that were tested for differential expression or differential splicing, respectively, in each dataset. For the differential expression analysis, the background genes are all that passed the minimum gene expression filter. For the differential splicing analysis, the background genes are limited to those genes for which rMATs found evidence of alternative splicing and were therefore tested for differential splicing between the marine and freshwater ecotypes. For the enrichment analysis of genes that were both differentially expressed and differentially spliced (DESGs), we used the intersection of the DEG and DSG background genes. To compare whether genes in EcoPeaks had stronger differential expression or differential splicing than genes outside EcoPeaks, we used a permutation test (10 000 permutations) to compare whether the median of the distribution of log2 fold-change of DEGs or the distribution of isoform difference of DSGs was different for genes inside and outside of EcoPeaks.

To test whether the DEGs and DSGs we identified might be involved in phenotypic divergence between the ecotypes, we did a similar enrichment analysis for quantitative trait loci (QTL) that underlie traits that differ between Pacific marine and freshwater populations (Peichel and Marques 2017; Rennison and Peichel 2022). We used the dataset of Liu et al. (2022), which excluded QTL that did not go in the expected direction (i.e QTL with marine alleles that lead to more freshwater-like phenotypes and vice-versa) (Liu et al. 2022). We lifted the coordinates of the QTL windows from v1 to v5 of the stickleback genome using the chain file “v1_withChrUn_to_v5.chain.txt” available at the Stickleback Genome Browser (https://stickleback.genetics.uga.edu/). Similar to the EcoPeak enrichment analysis, we: 1) identified the proportion of DEGs and DSGs that overlapped with QTL; 2) identified the proportion of background genes that overlapped with QTL; and 3) tested if the proportion of DEGs and DSGs in QTL is significantly higher than the proportion of background genes using a permutation test (1000 permutations) in R. We did this for the complete set of QTL and also for the different phenotypic categories of QTL defined by Peichel and Marques (2017): defence, behaviour and sensory system, body shape, body size, swimming, feeding, pigmentation, and respiration (Supplementary Table S4). We did not test for QTL category enrichment for the different types of DSGs because we did not have enough genes for the analysis to have statistical power.

### Effect of genetic distance in expression and splicing

To complement the EcoPeak enrichment analysis, we also looked for evidence of transcriptome-wide selection for stronger differential expression and splicing by looking at the correlation of splicing isoform difference and gene expression fold-change with genetic divergence between ecotypes. A significant positive correlation could mean transcriptome-wide selection for stronger differential expression (DE) and differential splicing (DS). To obtain a measure of genetic distance in both coding and non-coding regions across the stickleback genome, we used the SNP-level p-value data used to identify Pacific EcoPeaks by Kingman et al., 2021 (data kindly provided by the authors). These p-values result from a Fisher Exact Test for the probability of an imbalance in allele counts between ecotypes at each SNP (for more details consult Supplementary Section 9 in Kingman et al., 2021). A p-value < 0.05 means the SNP differs in allele frequencies between ecotypes and suggests divergent selection. As for the EcoPeak analysis, we translated all SNP coordinates from the v4.1 (*gasAcu1-4*) genome to the v5 coordinates. Then, for each gene in the transcriptome, we calculated a gene-level genetic divergence based on the average p-value of the SNPs overlapping each gene. With this data we tested if there was a correlation between genetic distance and strength of differential expression (using the log2fold-change metric from EdgeR) or splicing (measured as isoform difference, as described above) using all the genes tested in the differential expression and differential splicing analyses, separately. In the case of genes with more than one alternative splicing event, the isoform difference of the most differentially spliced event between ecotypes was selected. To test the significance of the correlations we used a non-parametric linear model based on ranks implemented in the *R* package *Rfit* v0.24.2 (Kloke and McKean 2012). We also did this analysis separately for genes inside and outside of EcoPeaks.

### Comparisons between types of AS

To see whether selection could be acting differently on particular types of AS, we compared both the genetic distance distributions and isoform difference distributions of the five types of AS events. DSGs were categorized into the five types of AS based on their most differential AS event. Then, we calculated the median of the isoform difference and genetic distance (average SNP p-value per gene) distributions for the five categories of DSGs, and we did pairwise tests on the difference of the medians using a Mann Whitney U Test, as implemented in the function wilcox.test() from the *R* package *stats* v3.6.1. In addition, we asked whether DSGs had a higher genetic distance than genes with alternative splicing but no significant differential splicing between ecotypes. For all genes with each type of AS event, we compared the median of the genetic distance distributions of the DSGs and the non-DSGs. We tested if this difference was significant with a permutation test (10 000 permutations).

### Gene Ontology enrichment analysis

We tested whether DSGs and DEGs were enriched in specific biological processes, through a Gene Ontology Enrichment analysis. We used the g:Profiler (Reimand et al. 2007) as implemented in the *R* package *gprofiler2* v0.2.2. We used the *G. aculeatus* functional annotations from Ensembl implemented in *gprofiler* and used custom backgrounds for the statistical analysis. As for the EcoPeak and QTL enrichment analysis, the background for the DEGs was the set of genes that passed the minimum expression, and the background for the DSGs was the set of genes with evidence of AS that passed the minimum expression filter junction filtering process (i.e the set of genes that were tested for DE by EdgeR and DS by rMATs). In addition, we also tested the enrichment of the subset of DEGs and DSGs that overlapped both Pacific EcoPeaks and QTL. Finally, we manually looked at the gene function of DSGs that overlapped both Global EcoPeaks and QTL in the GeneCards database (https://www.genecards.org/), which integrates information from several databases including NCBI Gene (https://www.ncbi.nlm.nih.gov/gene) and UniProt (https://www.uniprot.org/).

## Supporting information

Table S1 - RNAseq data used

Table S2 - DEGs and DSGs

Table S3 - GO analysis

Table S4 - QTL data

Table S5 - QTL enrichment analysis

Table S6 - EcoPeak enrichment results

Table S7 - DSGs overlapping Global EcoPeaks and QTL

Table S8 - EcoPeak and DS DE effect size

Table S9 - AS types comparison

Table S10 - DSGs vs non-DSGs genetic distance permutations

Table S11 - EcoPeaks

## Data availability

All RNAseq data used in this study was already publicly available at NCBI’s Sequence Read Archive (SRA) under BioProjects PRJNA371616 and PRJNA530695. Accession numbers for all samples used can be found in Supplementary Table S1. The EcoPeak data can be obtained from the UCSC Genome Browser (http://genome.ucsc.edu/) (instructions for downloading the data are in Kingman et al., 2021, under the “Data and materials availability” section). QTL data can be found in the supplementary material of Rennison and Peichel, 2022 and Liu et al., 2022. Scripts used for the analysis will be submitted to GitHub before publication.

## Acknowledgements

We are grateful to: Daniel Jeffries for very helpful discussion and feedback on the manuscript; Mike White for providing valuable chain files to lift the EcoPeaks from their v4.1 (*gasAcu*1-4) genomic coordinates to version 5 coordinates; Garrett Roberts Kingman, Krishna Veeramah and David Kingsley for kindly providing the SNP P-value data for our genetic distance analyses; Virginie Courtier-Orgogozo for kindly suggesting and helping with the search of phenotypes caused by splicing changes in GePheBase; and all Peichel Lab member for multiple helpful discussions during the realization of this study. This work was funded by the University of Bern.

