## Supplementary Figures for "The role of alternative splicing in marine-freshwater divergence in threespine stickleback"

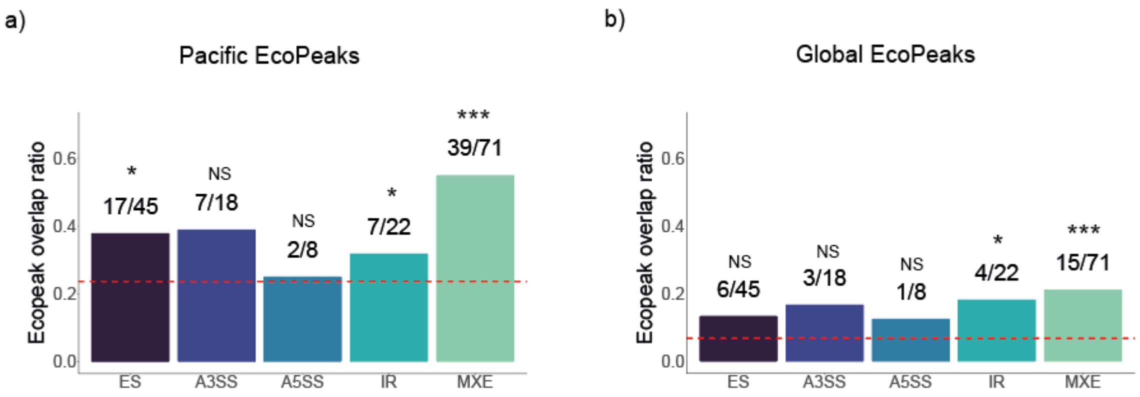


**Supplementary Figure S1**. Proportion of differential splicing events of each type found in genes within Pacific EcoPeaks (a) or Global EcoPeaks (b). Red dashed line represents the proportion of genes in the transcriptome that are found inside EcoPeaks. On top of each bar are shown the proportions of significantly divergent splicing events inside the EcoPeaks versus the total number of divergent splicing events of that type. Asterisks represent significance levels for the EcoPeak enrichment test (permutation test, 1000 permutations): * p-value < 0.05; ** p-value < 0.01; *** p-value < 0.001; NS – not significant, p-value > 0.05.

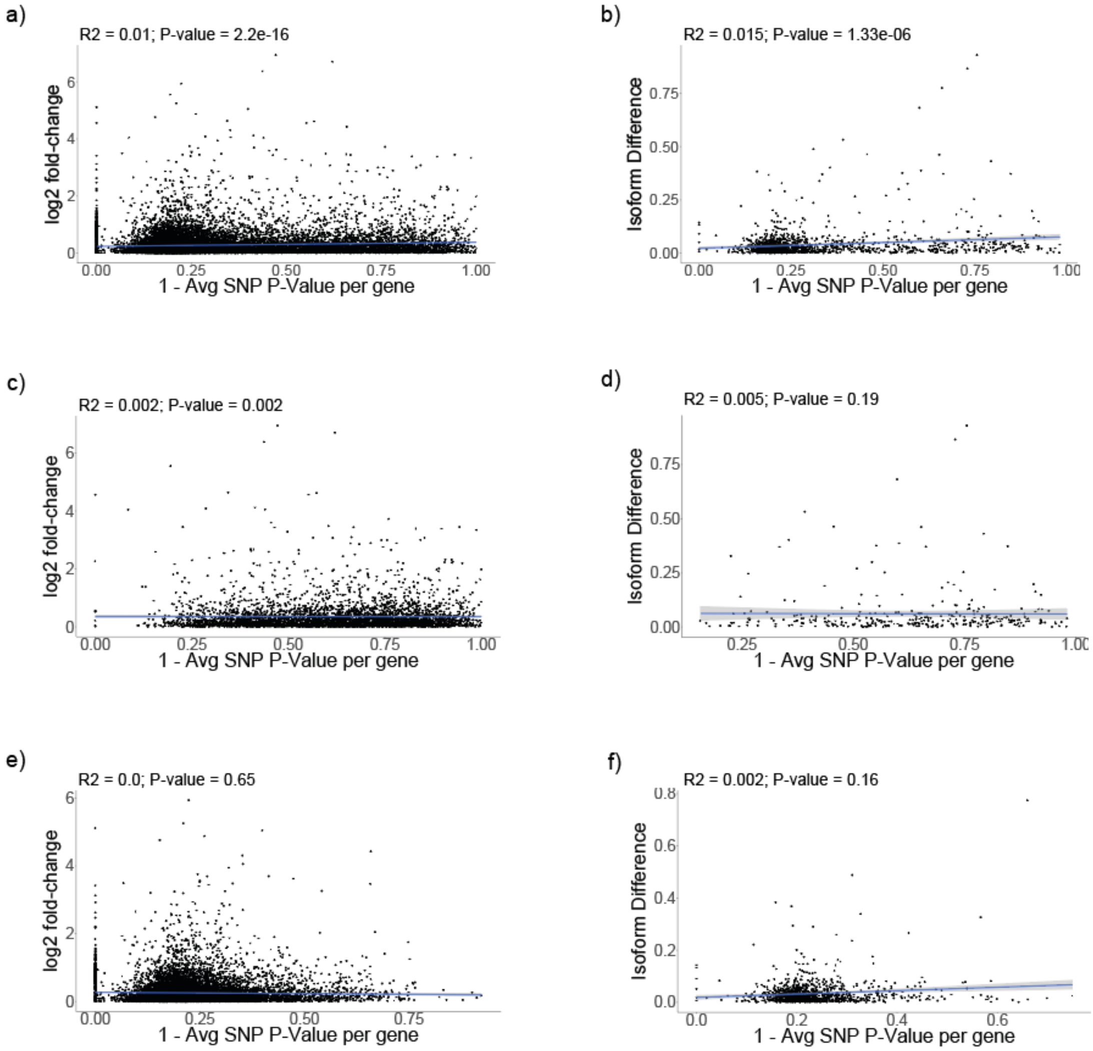


**Supplementary Figure S2.** Correlation between SNP average p-value and log2 fold-change (a, c, e) or isoform difference (b, d, f) for all background genes of the Pacific DE and DS analysis (a and b), only the background genes inside Pacific EcoPeaks (c and d), or only background genes outside of Pacific EcoPeaks (e and f). We subtracted the average SNP p-value to 1 so that higher values on the X axis represent higher genetic divergence of the SNPs in the genes.

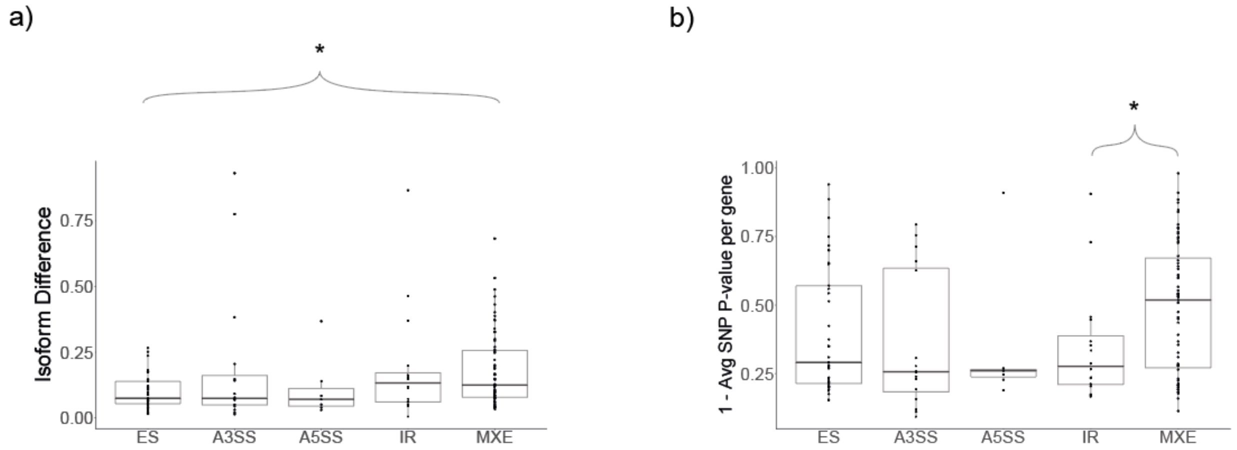


**Supplementary Figure S3.** Comparisons of the distribution of isoform differences (a) and SNP average p-value (b) between DSGs with different types of alternative splicing. Only the strongest splicing event per DSG was considered. In panel b) we subtracted the average SNP p-value from 1 so that larger values on the y-axis represent greater genetic divergence. Asterisks represent significant differences in the medians of the distributions according to a Mann Whitney U Test (p-value < 0.05). All pairwise combinations were tested, and only significant differences are shown.
